# Apparent sequential effects due to trial-to-trial fluctuations in decision criterion

**DOI:** 10.64898/2026.11.05.764575

**Authors:** Robin Vloeberghs, Kobe Desender, Anne E. Urai

## Abstract

Decision making is shaped by sequential effects, whereby preceding stimuli and responses influence current decisions. Computational models typically explain these effects by assuming that decision parameters are *causally* updated as a function of experimental history. Here, we challenge this assumption by showing that several hallmark sequential effects can instead emerge from slow fluctuations in the decision criterion. Using simulations based on Signal Detection Theory (SDT), we demonstrate that trial-to-trial criterion fluctuations, *independent* of experimental history, generate three commonly observed sequential effects: the repulsive effect of the previous stimulus (i.e., a tendency to alternate), the influence of previous response (i.e., choice history bias), and the modulation of choice history bias by previous confidence. To disentangle causal effects from effects driven by criterion fluctuations, we used the Hierarchical Model for Fluctuations in Criterion (hMFC), which simultaneously estimates both processes. Applying hMFC to 14 empirical datasets revealed that the repulsive effect of the previous stimulus was fully explained by criterion fluctuations. Accounting for criterion fluctuations also substantially altered choice history bias estimates: attractive biases disappeared in all four affected datasets, while three datasets revealed a previously masked tendency to alternate and two showed stronger alternation. In contrast, the modulation of choice history bias by previous confidence was largely unaffected by concurrent estimation of criterion fluctuations. Collectively, these findings demonstrate that criterion fluctuations account for a substantial portion of sequential effects and highlight the importance of explicitly modeling them to avoid confounded interpretations of behavioral and neuroimaging data.

## Introduction

Every day, we make numerous decisions, such as whether to drink coffee or tea, which T-shirt to wear, or whether to go to work by bike or car. In the laboratory, perceptual decision-making is often investigated using two-alternative forced-choice (2AFC) tasks, such as the random dot motion task or orientation discrimination tasks with Gabor patches. These paradigms provide a controlled framework for studying how observers transform sensory information into decisions. A prominent theoretical framework for understanding how decisions are formed is Signal Detection Theory (SDT) (Green & Swets, 1966). According to SDT, observers form an internal representation of task-relevant sensory information when presented with a stimulus (Figure 1A). This representation, often referred to as the decision variable, is then compared with an internal decision criterion to determine the decision. Because neural processing is inherently noisy, repeated presentation of the same stimulus does not produce identical decision variables. Instead, each stimulus is associated with a distribution of possible decision variables, resulting in variability in observers’ responses when the same stimulus is presented multiple times (Gold & Shadlen, 2007; Green & Swets, 1966; Macmillan & Creelman, 2005). In recent years, much work has focused on examining whether SDT models can be used to not just explain choices but also the sense of confidence that arises during decision-making. One influential account proposes that confidence is determined by the distance between the decision variable and the decision criterion, such that larger distances correspond with higher confidence (Hebart et al., 2016; Kepecs et al., 2008; Shekhar & Rahnev, 2021; Urai et al., 2017; Xue et al., 2024).

**Figure 1:**
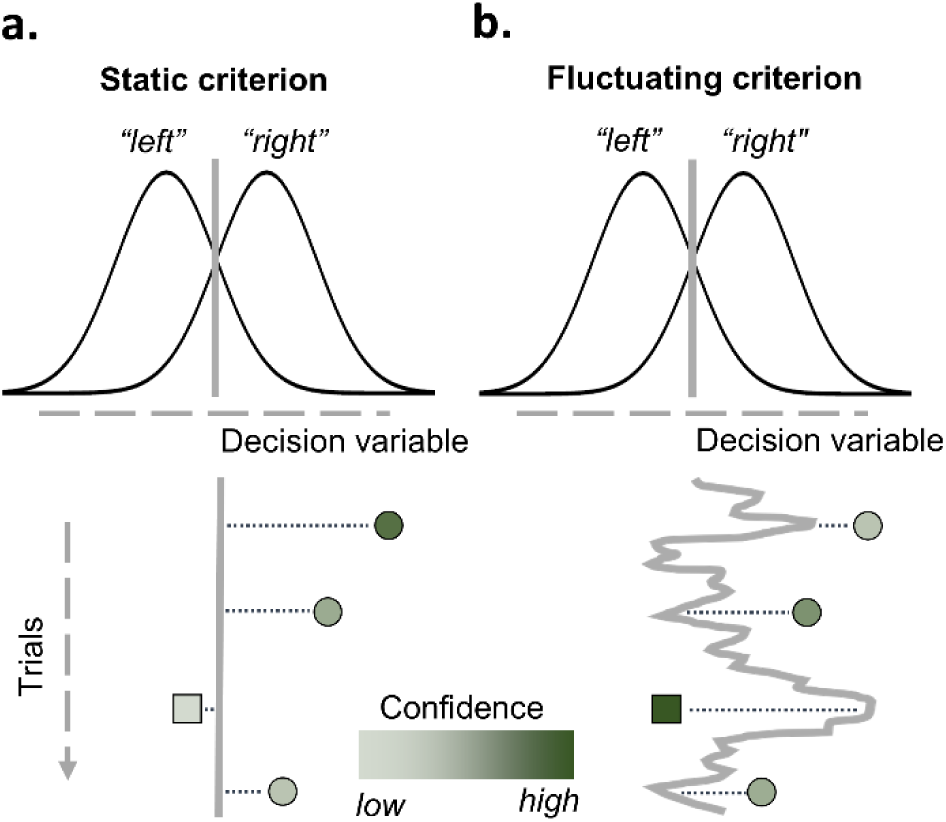
Signal Detection Theory (SDT) with static or fluctuating decision criterion. **A)** According to SDT, a choice is made by internally comparing a decision variable with a (static) decision criterion. If the decision variable exceeds the criterion, a “right” (circles) response will be made, else a “left” (squares) response will be made. Within this framework, confidence can be quantified as the absolute distance between the decision variable and the decision criterion, with a larger distance corresponding to higher confidence. **B)** While the decision criterion is typically assumed to be static over trials, increasing empirical work suggests that instead of being fixed the criterion tends to fluctuate from trial-to-trial (Cowley et al., 2020; Gupta & Brody, 2022; Roy et al., 2021; Vloeberghs, Navarrete, et al., 2025; Vloeberghs, Urai, et al., 2025).

In its simplest form, computational models such as Signal Detection Theory (SDT) describe how momentary sensory evidence is transformed into a perceptual decision and an accompanying sense of confidence. However, perceptual decisions do not occur in isolation; rather, they are embedded within a continuous sequence of experiences. A large body of research, in humans, primates, and rodents has shown sequential effects in decision-making: decisions are influenced not only by the current sensory evidence, but also by information from previous trials. Sequential effects have been observed in humans (Bosch et al., 2020; Braun et al., 2018; Fischer & Whitney, 2014; Fritsche et al., 2020; Fründ et al., 2014; Samaha et al., 2019; Schlunegger & Mast, 2023; Urai et al., 2017, 2019; Urai & Donner, 2022; Van Geert et al., 2022; Yu & Cohen, 2008), primates (Lueckmann et al., 2018; Mochol et al., 2021), and rodents (Fernandez et al., 2025; Gupta & Brody, 2022; Lak et al., 2020) for a range of tasks. Three specific observations have been made repeatedly: first, the current choice is influenced by the preceding choice with a tendency to repeat (or alternate) the previous response, an effect referred to as choice history bias, serial dependence, repetition bias, or hysteresis. Second, this influence of the previous response is additionally modulated by confidence with an increased repetition tendency following high confidence trials, and a tendency to alternate following (perceived) error trials (Bosch et al., 2020; Braun et al., 2018; de Gee et al., 2020; Samaha et al., 2019). Third, decisions are biased away from the previous stimulus direction, with a stronger effect for increasing stimulus strength (Bosch et al., 2020; Fritsche et al., 2017, 2020; Pascucci et al., 2019; Sadil et al., 2024; Van Geert et al., 2022).

To account for these sequential effects, previous work has generally assumed that they arise from the *causal* influence of experimental history, such as preceding responses or stimuli. Within Signal Detection Theory (SDT), for example, Treisman & Williams (1984) proposed that choice history biases emerge because the decision criterion is updated on each trial as a function of the previous response (see also Lages & Treisman, 2010). Likewise, criterion shifts have been used to explain repulsive effects of the previous stimulus, resulting in a tendency to alternate choices (Dyjas et al., 2012; Hachen et al., 2020; Lee et al., 2023; Lee & Lee, 2023). Other accounts have attributed choice history biases to reinforcement learning mechanisms that update action values based on previous responses (Akaishi et al., 2014), or to Bayesian ideal observer models in which preceding choices shape decision making through prior beliefs (Fritsche et al., 2020; Van Geert et al., 2024; Wei & Stocker, 2015).

Although these modeling approaches differ in their computational implementation, they share the assumption that experimental history *causally* influences subsequent decisions. However, this assumption may not be necessary. Growing evidence indicates that computational variables, like the decision criterion, tend to fluctuate over time (Figure 1B), presumably due to changes in attention, arousal, motivation, or other internal states (Cowley et al., 2020; Gupta & Brody, 2022; Roy et al., 2021; Schurr et al., 2024; Vloeberghs, Navarrete, et al., 2025; Vloeberghs, Urai, et al., 2025). If these fluctuations are sufficiently slow (i.e., autocorrelated), they can create temporal dependencies in behavior that resemble sequential effects, even without any causal influence of previous trials. Consistent with this idea, Vloeberghs and colleagues (2025) showed that allowing the SDT decision criterion to fluctuate across trials according to a first-order autoregressive (AR1) process is sufficient to generate choice history bias. Crucially, unlike previous work in which criterion updates are driven by the preceding response (Lages & Treisman, 2010; Treisman & Williams, 1984), these criterion fluctuations were generated *independently* of previous stimuli or responses (see also Gupta & Brody, 2022; Lak et al., 2020 for similar simulations). Intuitively, when the decision criterion gradually fluctuates to the right, leftward responses become more likely over a sequence of trials. As the criterion later fluctuates towards the left, rightward responses become more likely. These slow criterion fluctuations therefore induce autocorrelated response sequences (e.g., left, left, right, right), even in the absence of any true influence of previous responses.

Whereas previous work has shown that choice history bias can be explained either by history-dependent criterion updates or by an independently fluctuating decision criterion, the relative contribution of these two mechanisms remains unknown. Therefore, in the current work we will investigate to what extend sequential effects are driven by criterion fluctuations. Previous work using simulations with criterion fluctuations has focused exclusively on choice history bias. Therefore, we will use simulations to investigate whether criterion fluctuations can also account for other sequential effects, including the confidence-modulation of choice history bias and the repulsive effect of the previous stimulus. We will then use the Hierarchical Model for Fluctuations in Criterion (hMFC) (Vloeberghs, Urai, et al., 2025), which allows to simultaneously estimate criterion fluctuations and the causal influence of experimental history, to disentangle the contribution of each mechanism to sequential effects in 14 empirical datasets.

## Methods

### Empirical data

We selected empirical datasets based on the following inclusion criteria: (1) participants provided trial-by-trial confidence ratings using a confidence scale, (2) each participant completed at least 500 trials (i.e., to ensure robust estimation of trial-by-trial criterion with hMFC), 3) the main experimental manipulation was the strength of the stimulus presented on each trial; (4) no more than two stimuli were presented simultaneously (i.e., excluding tasks such as 2IFC paradigms with six Gabor patches); and (5) no trial-by-trial performance feedback was provided. Based on these criteria, we selected 12 datasets from the Confidence Database (Rahnev et al., 2020). We additionally included datasets from Boldt et al., (2017) and Vloeberghs et al. (unpublished), resulting in a total of 14 datasets.

The fourteen studies included a diverse range of perceptual decision-making tasks. Five studies used Gabor patches as stimuli, and four studies employed the random dot motion task, in which participants indicated whether they perceived leftward or rightward motion within a cloud of moving dots. Two studies required participants to judge which of two panels contained more dots. The remaining studies included an ambiguous face classification task (fearful vs. happy), a source-classification task involving a series of dots, and a task in which participants determined the average color of a group of shapes. The majority of the studies, 9 out of 14, used a discrete 4-point confidence scale, whereas the remaining five studies either used a continuous confidence ratings or a 2-, 3-, or 6-point scale. Sample sizes ranged from 10 to 81 participants, and the number of trials per study varied between 504 and 3240. Thirteen subjects were excluded because they either performed at chance level or provided the same confidence rating on over 90% of trials. An additional six subjects were removed due to estimated criterion states larger than 10. Values this large saturate the sigmoid-transformed log-odds in hMFC, potentially compromising the accuracy of the model estimates. After applying these exclusion criteria, the final dataset consisted of 498 participants and 484.962 trials. Additional details for each study are provided in Supplementary Table 1.

### Hierarchical Model for Fluctuations in Criterion

To quantify the relative contribution of experimental history (i.e., causal influence) and criterion fluctuations to sequential effects in empirical data, we relied on the Hierarchical Model for Fluctuations in Criterion (hMFC). This model predicts the binary response *y_t_* (0 or 1) at trial *t* using a Bernoulli distribution:

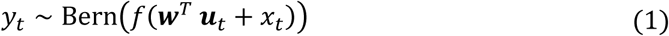

where the probability of a response is determined by a linear combination of the observed input variables *u_t_*, weighted by the parameter vector *w*, and a latent state *x_t_* representing the decision criterion on trial *t*. The sigmoid transformation *f*(*z*) constrains the probability between 0 and 1. The latent criterion evolves from trial-to-trial according to a first-order autoregressive (AR1) process with intercept *b*, autoregressive coefficient *ϕ*, and innovation variance *σ*^2^:

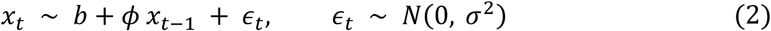

Adding the latent state *x_t_* to the Bernoulli log-odds is equivalent to modeling decisions as the comparison of a decision variable against a trial-by-trial fluctuating decision criterion. Note that the criterion state estimated by hMFC is expressed as a bias on the probability of making a particular response (e.g., a rightward response). Consequently, positive values of *x_t_* increase the probability of a right response, which corresponds to a leftward shift of the decision criterion in SDT. For ease of interpretation we have changed the sign of the hMFC output such that the interpretation of *x_t_* is similar to the SDT criterion.

The parameters *w*, *b*, *a* and *σ*^2^ are estimated per subject. In addition to estimating these subject-specific parameters, hMFC infers hierarchical (group-level) distributions that allow information to be shared across subjects, improving the estimation accuracy of individual parameter estimates. Through this hierarchical structure, hMFC can accurately recover the model parameters, even with as few as 500 trials per subject. Most importantly, hMFC demonstrates an excellent recovery of the latent decision criterion trajectories, providing reliable trial-to-trial estimates of the decision criterion. A detailed description of the model and its parameter recovery is provided in Vloeberghs, Urai, et al., (2025).

In the current study, as input variables ***u****_t_* we included the current stimulus, previous stimulus, previous response, previous confidence, the interaction between previous response and previous confidence. All input variables were scaled to range between −1 and +1. Importantly, including these predictors allows the model to estimate sequential effects while controlling for the influence of latent criterion fluctuations.

The 14 empirical datasets were each be fitted with hMFC both with and without estimating criterion fluctuations. The extent to which criterion fluctuations drive sequential effects were quantified by directly comparing posterior distributions of the two models. For the analysis *across* all datasets, we computed an averaged posterior distribution over all datasets. Specifically, for each posterior draw, we averaged the corresponding samples across all datasets, resulting in an aggregated posterior distribution that contains the same number of posterior samples as the individual dataset-level posteriors. The influence of criterion fluctuations on sequential effects was additionally summarized by using the reduction metric:

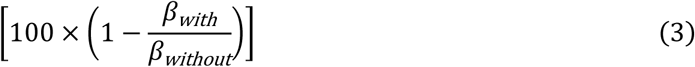

where *β_with_* and *β_without_* denote the posterior means estimated with and without criterion fluctuations, respectively. A value of 0% indicates that accounting for criterion fluctuations does not change the estimated effect, whereas a value of 100% indicates that the effect is completely eliminated. Values greater than 100% indicate that the estimated effect reverses sign after accounting for criterion fluctuations (e.g., from positive to negative), while negative values indicate that the estimated effect increases in magnitude. To avoid interpreting percentage changes in effects that were absent, the reduction metric was only calculated when the 95% credible interval of at least one of the two posterior estimates excluded zero.

### Simulations

To investigate whether sequential effects can be generated under the mere presence of criterion fluctuations we simulated 5000 trials from a SDT observer (Figure 1B). On each trial, a decision variable was sampled from one of two normal distributions with mean *μ* = [-.50, -.25, .25, .50] and standard deviation *σ* = 1. The decision criterion varied across trials according to a first-order autoregressive process, AR(1), with autoregressive coefficient *ϕ* = .9995 and innovation standard deviation *σ* = .01. Crucially, these criterion fluctuations occurred fully independent of experimental history such as previous response or previous stimulus, therefore no causal influence was present. A binary response was generated on each trial by comparing the decision variable with the decision criterion. Confidence was computed using a post-decisional evidence sample (Fleming et al., 2018). Following the initial decision, an additional evidence sample was drawn from the same generative distribution as the pre-decisional evidence. Confidence was then computed by combining the signed distances of both the pre- and post-decisional evidence samples from the decision criterion. When both evidence samples supported the chosen response, confidence increased. Conversely, when the post-decisional evidence favored the unchosen alternative, confidence decreased. This approach allows the simulation of perceived error trials, something that traditional SDT models grapple with (Shekhar & Rahnev, 2024). This allows us to investigate the full confidence-dependent modulation of choice history bias, including increased repetition following high-confidence decisions and increased alternation following perceived errors.

To determine whether sequential effects emerged in the simulated data, we fit a generalized linear model predicting binary responses from the current stimulus (i.e., the mean of the generative normal distribution), the previous stimulus, the previous response, the interaction between previous response and previous confidence:

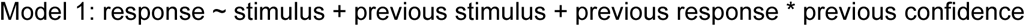

We then fit a second model to test the central premise of hMFC: if apparent sequential effects are driven by latent criterion fluctuations, explicitly accounting for these fluctuations should eliminate the sequential effects, causing the corresponding regression coefficients to become non-significant:

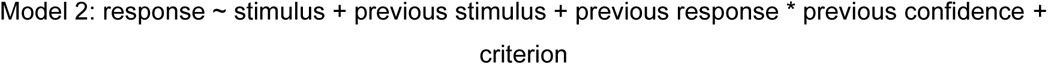

Together, these simulations allow us to evaluate whether latent criterion fluctuations alone can generate apparent sequential effects and whether including the criterion in the model accounts for these effects.

## Results

### Simulation study

To investigate the role of criterion fluctuations in sequential effects we simulated 5000 trials for an SDT observer with the criterion fluctuating from trial-to-trial according to an AR(1) process (Figure 1B). Importantly, the simulation contained no causal influences of experimental history: the decision criterion was not affected by previous stimulus, previous responses, or previous confidence. Nevertheless, predicting the current response using a generalized linear model revealed the presence of sequential effects. First, there was a significant effect of previous stimulus (*β* = −4.196, SE = 0.323, *χ*^2^(1) = 179.34, *p* < .001, CI(95) = [-4.989, −3.562]), indicating an increased tendency to alternate as the stimulus strength increases (Figure 2A). Second, we found a choice history bias and the modulation by confidence as the interaction between previous response and previous confidence was significant (*β* = 6.269, SE = 0.232, *χ*^2^(1) = 1052.30, *p* < .001, CI(95) = [5.822, 6.73]). When confidence on the previous trial was high, a tendency to repeat that response was observed, whereas low confidence (i.e., a likely error) promotes a tendency to alternate (Figure 2B). The main effect of previous response was not significant (*β* = 0.019, SE = 0.058, *χ*^2^(1) = 0.11, *p* = .740, CI(95) = [-0.094, 0.132]), indicating that there was no evidence for a uniform choice history bias across levels of previous confidence (i.e., no overall tendency to repeat or alternate). Note that when previous confidence was omitted as predictor, there was a significant attractive effect of previous response. Without the inclusion of previous confidence, the main effect of previous response reflects the average tendency to repeat or alternate across all trials. Since confidence ratings were, on average, above zero (mean = 2.716), the stronger repetition tendency following high-confidence trials dominates this average, resulting in an overall attractive effect of previous response. Lastly, the effect of stimulus was significant (*β* = 4.487, SE = 0.319, *χ*^2^(1) = 214.53, *p* < .001, CI(95) = [3.867, 5.117]), whereas the main effect of previous confidence was absent (*β* = −0.33, SE = 0.22, *χ*^2^(1) = 2.27, *p* = .132, CI(95) = [-0.762, 0.099]). No multicollinearity issues were present (all VIFs < 2.286). Together, these simulations demonstrate that criterion fluctuations, despite being generated independently of experimental history, are sufficient to produce sequential effects.

**Figure 2:**
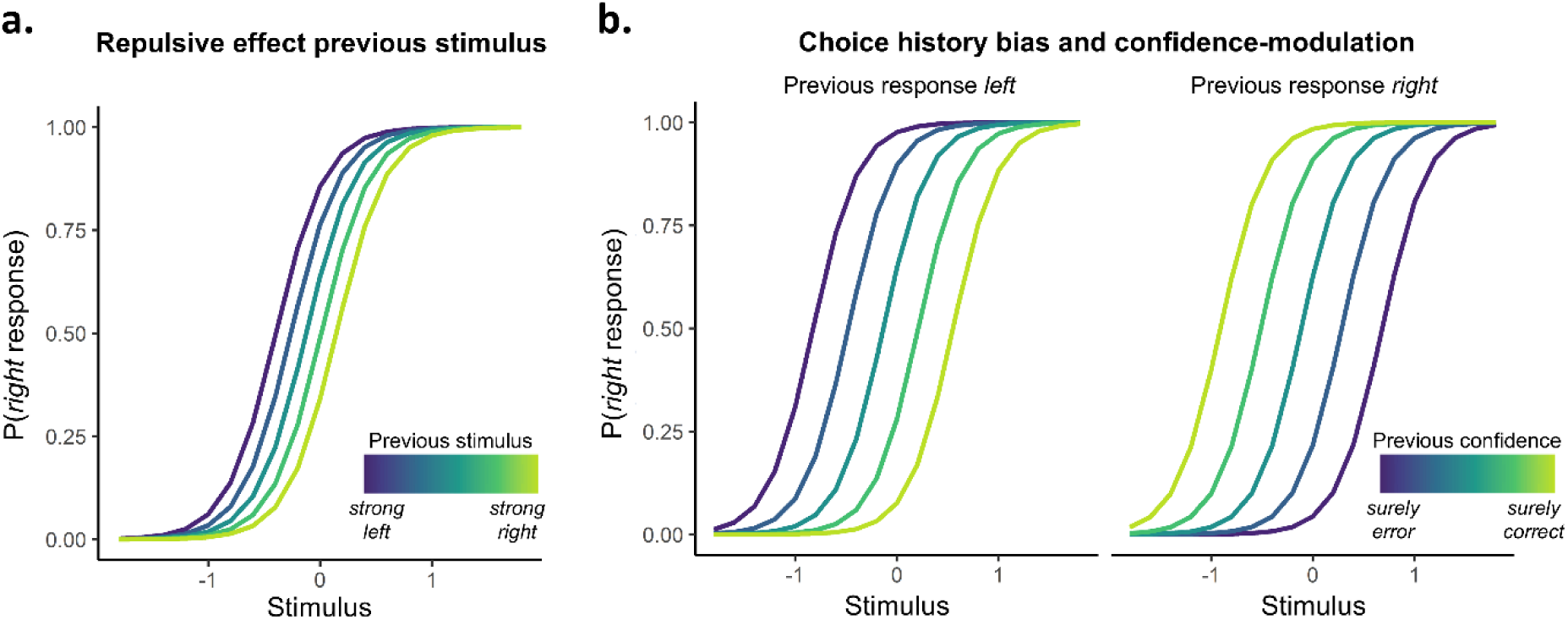
Criterion fluctuations generate sequential effects in simulated data. Data simulated from an SDT observer (5000 trials) with the criterion fluctuating across trials according to an AR(1) process (Figure 1B). **A.** Although the criterion was not causally influenced by experimental history, an apparent repulsive effect of the previous stimulus emerged, with an increasing tendency to alternate responses as the strength of the previous stimulus increased. **B.** In addition, the simulated data exhibited a choice history bias that was modulated by confidence on the previous trial. Specifically, following a left response made with low confidence (i.e., surely error), observers were more likely to make a right response on the current trial. In contrast, following a left response made with high confidence, observers were less likely to make a right response. More generally, high confidence on the previous trial promoted a repetition of the previous response, whereas low confidence promoted response alternation.

Next, we tested the central premise of hMFC: that if sequential effects arise from trial-to-trial fluctuations in the decision criterion, explicitly accounting for these fluctuations should eliminate the apparent history effects in simulated data (see also Figure 6 in Vloeberghs, Urai, et al. (2025) showing that hMFC controls for apparent choice history bias while accurately recovering the true effect of previous response). As a proof of concept, we added the generative trial-by-trial criterion values to the generalized linear model. Whereas the model without criterion estimates showed a significant main effect of previous stimulus and a significant interaction between previous response and previous confidence, both effects disappeared after including the generative criterion fluctuations (previous stimulus: *p* = 0.893; previous response × previous confidence: *p* = 0.935). The main effect of criterion was significant (*β* = −1.805, SE = 0.079, *χ*^2^(1) = 711.57, *p* < .001, CI(95) = [-1.961, −1.653]), indicating an increased probability of making a right response when the criterion fluctuates towards the left. Also the effect of stimulus was significant (*β* = 5.66, SE = 0.373, *χ*^2^(1) = 262.51, *p* < .001, CI(95) = [4.937, 6.398]). Lastly, the effect of previous response (*p* = 0.497) and previous confidence (*p* = 0.805) were both non-significant. No multicollinearity issues were present (all VIFs < 4.565). Together, these simulations demonstrate that criterion fluctuations alone can give rise to apparent sequential effects, including the confidence-modulation of choice history bias and the repulsive influence of the previous stimulus. Importantly, once these criterion fluctuations are explicitly accounted for, these apparent sequential effects disappear.

### Empirical data

Following these simulations, we next investigated the extent to which sequential effects can be explained by trial-to-trial fluctuations in the decision criterion in empirical data. To this end, we estimated trial-by-trial criterion fluctuations using hMFC (Vloeberghs, Urai, et al., 2025). In simulated data, hMFC accurately recovers latent criterion fluctuations, even with as few as 500 trials per participant (Figure 3A-C). In empirical data, the estimated criterion trajectories capture systematic changes in response probabilities that are consistent with shifts in the decision criterion predicted by SDT (see Figure 3D for an example dataset). Specifically, rightward shifts in the estimated criterion are associated with an increased probability of leftward responses, whereas leftward shifts are associated with an increased probability of rightward responses. Model fitting produced stable and well-behaved posterior distributions, with good mixing across Gibbs sampler chains (all R-hat < 1.05), indicating reliable sampling from the posterior distribution. Together, these findings provide strong evidence that hMFC yields valid and reliable estimates of trial-by-trial criterion fluctuations.

**Figure 3:**
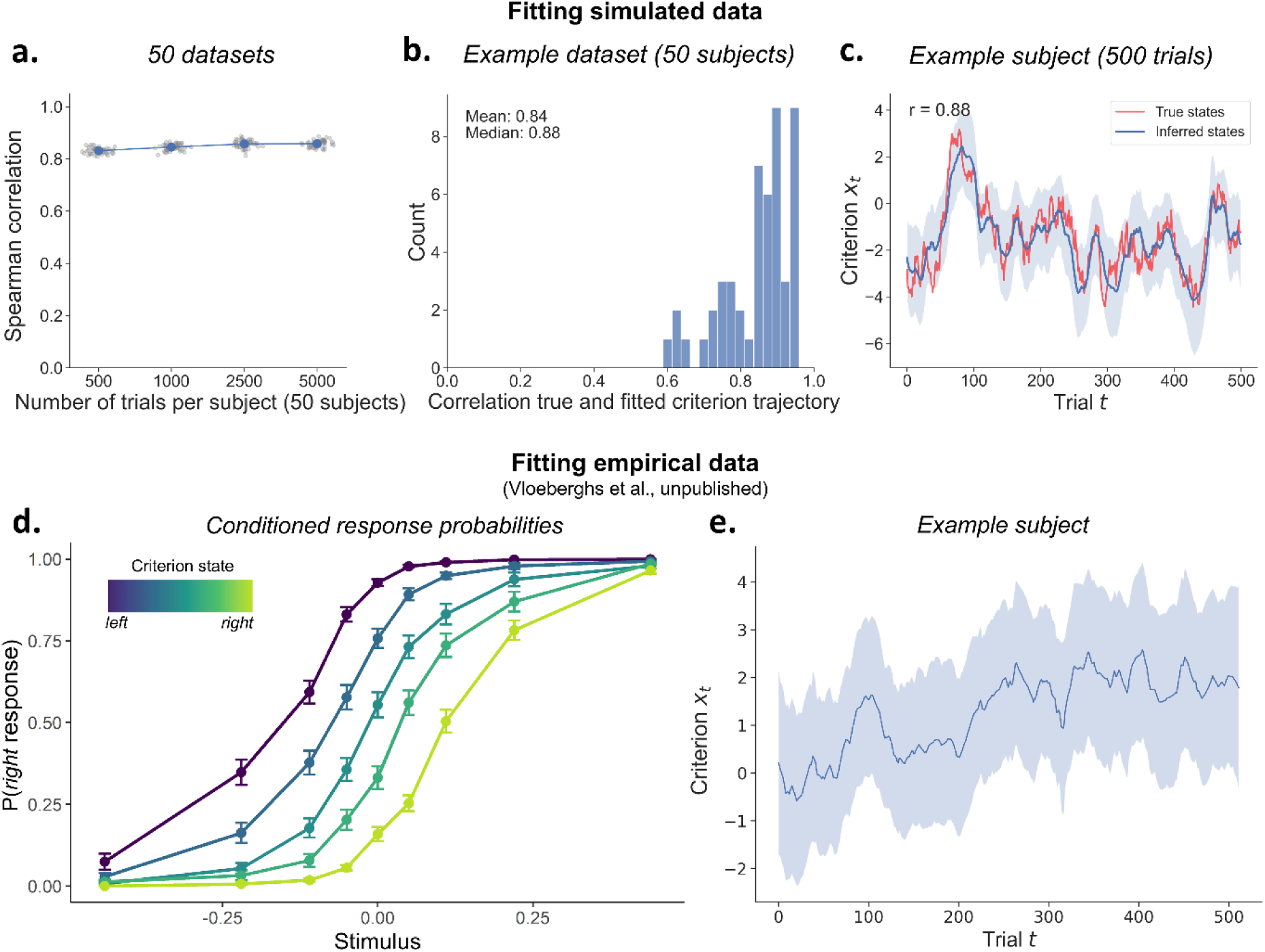
Using hMFC to estimate criterion fluctuations in simulated and empirical data. **A.** The recoverability of trial-by-trial criterion fluctuations by hMFC, as evaluated in Vloeberghs et al. (2025) using simulated datasets containing 50 subjects and varying numbers of trials per subject (500, 1000, 2500, or 5000; 50 datasets per trial count). Each point represents the average correlation between the true and estimated criterion trajectories across subjects within a single dataset. Error bars denote the standard error of the mean but are barely visible due to their small size. **B.** Correlations between the true and estimated criterion trajectories for individual subjects in a representative simulated dataset with 500 trials per subject. **C.** True and estimated criterion trajectories for an example simulated subject with 500 trials. The shaded region indicates the 95% posterior credible interval at each trial. **D.** Empirical response probabilities from Vloeberghs et al. (unpublished), conditioned on binned trial-by-trial criterion estimates. As the criterion shifts towards the left, the probability of a rightward response increases. **E.** Estimated criterion trajectory for an example participant from the same dataset, with the shaded region representing the 95% posterior credible interval.

#### Effect of previous stimulus

Having established that hMFC provides reliable estimates of criterion fluctuations, we next tested whether these fluctuations account for sequential effects in empirical data. To this end, we fitted each dataset using hMFC once with and once without estimating trial-by-trial criterion fluctuations. If sequential effects are entirely driven by criterion fluctuations, then accounting for these fluctuations should eliminate the estimated sequential effects. We first focused on the finding that the strength of the *previous stimulus* tends to be associated with an increased tendency to alternate responses. Across all datasets, we observed a significant repulsive effect of the previous stimulus when criterion fluctuations were not estimated (*β_without_* = −0.497, CI(95) = [-0.665, −0.329]), with *β_without_* representing the posterior mean. This effect indicates an increased tendency to alternate responses as the strength of the previous stimulus increased (Figure 4A, *Total*). Critically, when fitting the same model to the data but while additionally estimating criterion fluctuations, this effect was substantially reduced and was no longer credibly different from zero (*β_with_* = −0.113, CI(95) = [-0.258, 0.03]), indicating that the repulsive effect of previous stimulus is entirely driven by criterion fluctuations.

**Figure 4:**
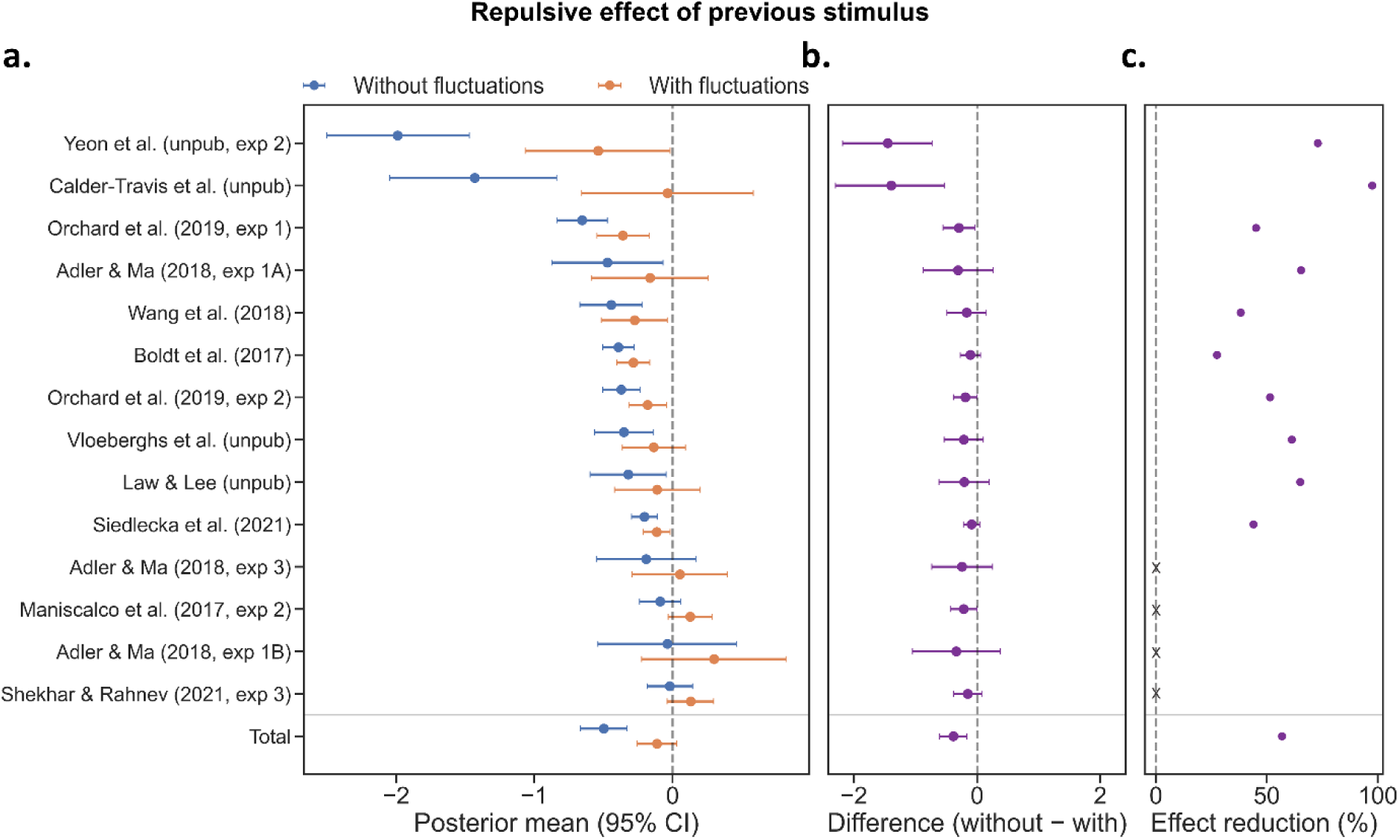
The effect of previous stimulus disappears after controlling for criterion fluctuations. **A.** Posterior means and 95% credible interval for the previous stimulus coefficient estimated with hMFC, with and without criterion fluctuations. Without estimating criterion fluctuations, a significant repulsive effect of the previous stimulus (i.e., a tendency to alternate responses) was observed in 10 of 14 datasets. After accounting for criterion fluctuations, this effect remained significant in only 6 of 14 datasets. The pooled posterior estimate across all datasets (bottom; Total) showed the same pattern: the significant previous stimulus effect disappeared once criterion fluctuations were explicitly modeled. **B.** The estimated previous stimulus effect was smaller after accounting for criterion fluctuations in every dataset (i.e., all posterior differences were negative), although this reduction reached statistical significance in only 5 of the 14 datasets. The pooled analysis nevertheless revealed a significant overall reduction in the previous stimulus effect after controlling for criterion fluctuations. **C.** Across datasets, the estimated previous stimulus effect was reduced by an average of 56.8% (range: 27.5 - 97.5%) after including criterion fluctuations. To avoid interpreting percentage changes for effects that were not reliably present, this reduction metric was computed only for datasets in which the 95% credible interval of at least one of the two posterior estimates excluded zero.

We then examined the datasets individually. Without accounting for criterion fluctuations, a significant repulsive effect of the previous stimulus was observed in 10 of the 14 datasets. After including criterion fluctuations, this effect was no longer supported in 4 of these 10 datasets, although it remained significant in the other 6. Nevertheless, comparing the posterior estimates obtained with and without criterion fluctuations (Figure 4B) revealed a highly consistent reduction in the previous stimulus effect in all 14 datasets (i.e., negative posterior differences), with the 95% credible interval excluding zero in 5 datasets. Consistent with this pattern, the pooled posterior difference across datasets was also credibly below zero (*β_without−with_* = −0.384, CI(95) = [-0.606, −0.165]).

Finally, among the datasets showing a significant previous stimulus effect before accounting for criterion fluctuations, the estimated effect was reduced by an average of 56.8% (range: 27.5 - 97.5%) after criterion fluctuations were included (Figure 4C). Together, these findings indicate that trial-by-trial criterion fluctuations entirely explain the repulsive effect of the previous stimulus.

#### Effect of previous response

Second, we investigated the influence of the *previous response* on the current response, referred to as choice history bias. When considering all datasets combined, no choice history bias was found when criterion fluctuations were not estimated (*β_without_* = 0.073, CI(95) = [-0.054, 0.199]) (Figure 5A, *Total*). Also after explicitly controlling for criterion fluctuations, the effect of previous response was not significant (*β_with_* = −0.108, CI(95) = [-0.223, 0.008]). Closer inspection of the individual datasets, however, revealed substantial heterogeneity in choice history bias (Figure 5A). Specifically, when not accounting for criterion fluctuations 4 of the 14 datasets showed an attractive bias, reflecting a tendency to repeat the previous response (i.e., positive posterior means), whereas 2 datasets showed a repulsive bias, reflecting a tendency to alternate (i.e., negative posterior means). Interestingly, accounting for trial-by-trial criterion fluctuations eliminated the attractive previous response bias in all four datasets, whereas the repulsive bias observed in the two datasets remained. Moreover, in three additional datasets, controlling for criterion fluctuations uncovered a significant tendency to alternate that was not present in the model without criterion fluctuations. Thus, criterion fluctuations appear to substantially influence estimates of choice history bias. Accounting for these fluctuations systematically shifts these estimates toward alternation.

**Figure 5:**
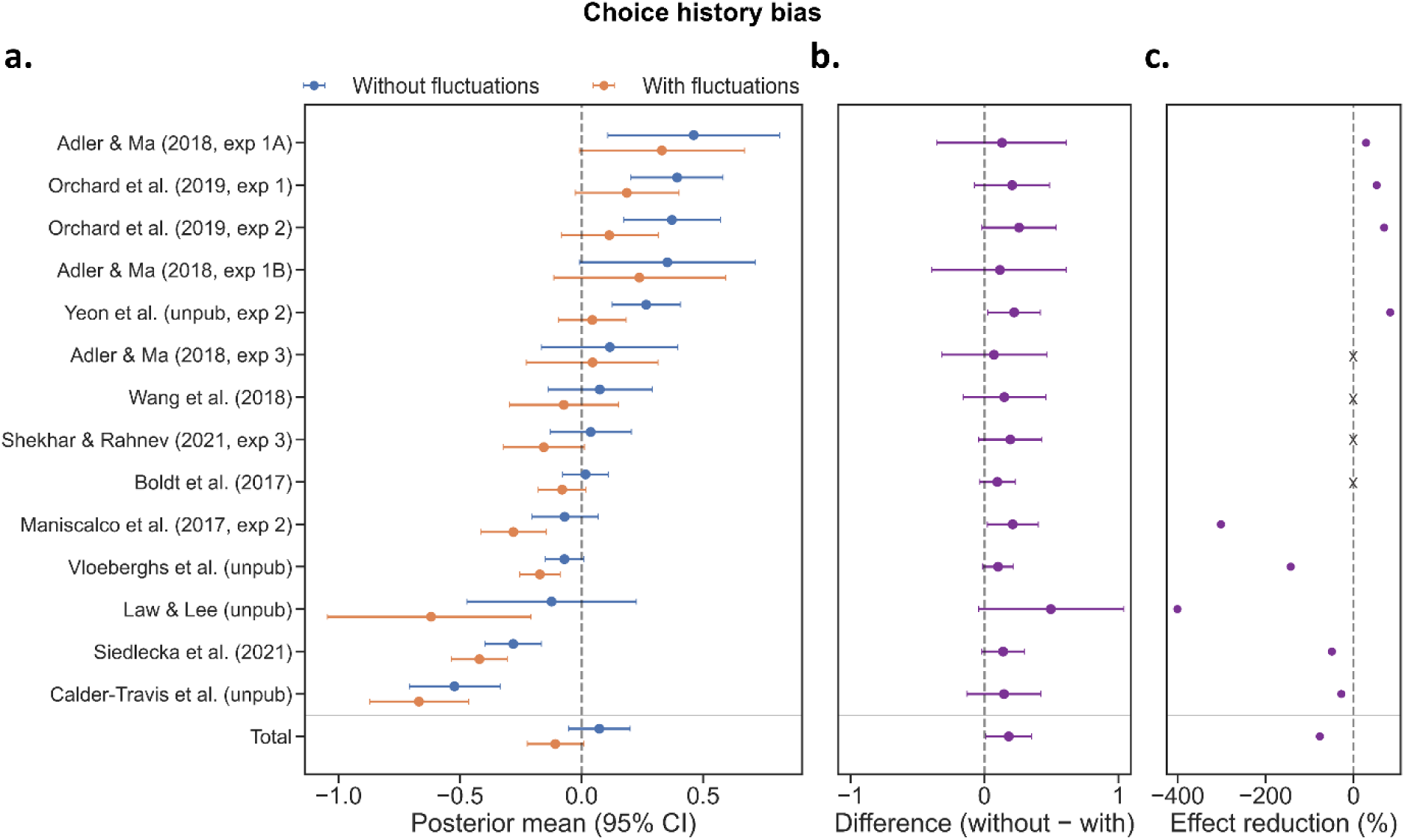
Accounting for criterion fluctuations steers choice history bias towards alternation. **A.** Posterior means and 95% credible interval for the previous response coefficient estimated with hMFC, with and without criterion fluctuations. Without accounting for criterion fluctuations, 4 of the 14 datasets exhibited a significant attractive effect of the previous response. After controlling for criterion fluctuations, this effect disappeared in all four datasets. In three additional datasets, accounting for criterion fluctuations revealed a significant repulsive effect (i.e., a tendency to alternate) that was absent in the model without criterion fluctuations. In two datasets that already showed a repulsive effect, the magnitude of this effect increased after criterion fluctuations were included. Across all datasets combined (bottom; Total), no significant previous response effect was observed in either model. **B.** Posterior differences between the two models indicate that accounting for criterion fluctuations systematically shifts choice history bias toward alternation, with attractive effects diminishing and repulsive effects becoming stronger. At the individual dataset level, these differences were significant in 2 of the 14 datasets, whereas the effect was significant across all datasets combined. **C.** In the four datasets where the attractive effect disappeared after accounting for criterion fluctuations, the estimated previous response effect was reduced by an average of 58.5%. Across the remaining datasets, accounting for criterion fluctuations increased the magnitude of the repulsive effect by an average of 184.4% (negative percentage values in the plot correspond to a stronger tendency to alternate). To avoid interpreting percentage changes for effects that were not reliably present, this reduction metric was computed only for datasets in which the 95% credible interval of at least one of the two posterior estimates excluded zero.

Strikingly, the posterior differences shown in Figure 5B demonstrate that in all studies considered, accounting for criterion fluctuations shifted the estimated previous response bias towards alternation: attractive biases became weaker, whereas repulsive biases became stronger (*β_without−with_* = 0.181, CI(95) = [0.009, 0.353]). Interestingly, although this effect was highly consistent across studies, it was only statistically significant in 2 of the 14 datasets.

Finally, the four datasets in which the attractive effect of the previous response disappeared after accounting for criterion fluctuations showed an average reduction of 58.5% (range: 28.4 - 83.4%) (Figure 5C). In contrast, the three datasets in which controlling for criterion fluctuations revealed a significant repulsive effect exhibited an average increase of 281.7% (range: −400.6 to −142.8%). These seemingly large percentage changes arise because the posterior means before accounting for criterion fluctuations were close to zero, making the relative change highly sensitive (see Methods for details of the computation). In the two datasets where the repulsive effect was already present, accounting for criterion fluctuations further strengthened it by an average of 38.6%. Across all datasets, accounting for criterion fluctuations shifted the estimated choice history bias toward a more repulsive effect, with an average change of 76.5% (median: 27.8%). Together, these findings demonstrate that trial-by-trial criterion fluctuations substantially influence estimates of choice history bias.

#### Effect of previous response x previous confidence

Third, we investigated whether criterion fluctuations influence the *confidence modulation of choice history bias* (i.e., the interaction between previous response and previous confidence). When criterion fluctuations were not accounted for, we observed a significant effect across all datasets (*β_without_* = 0.215, CI(95) = [0.06, 0.372]), showing that participants were more likely to repeat choices when these were made with high confidence, but alternate choices when these were perceived to be erroneous (Figure 6A, *Total*). Interestingly, contrary to the previously discussed two effects, this effect was almost entirely unaffected by incorporating criterion fluctuations into the model and the effect remained significant (*β_with_* = 0.179, CI(95) = [0.04, 0.321]). At the individual dataset level, a significant confidence-modulation of choice history bias was observed in 4 out of 14 datasets. All four effects remained significant after controlling for criterion fluctuations, while the effect emerged in two additional datasets.

**Figure 6:**
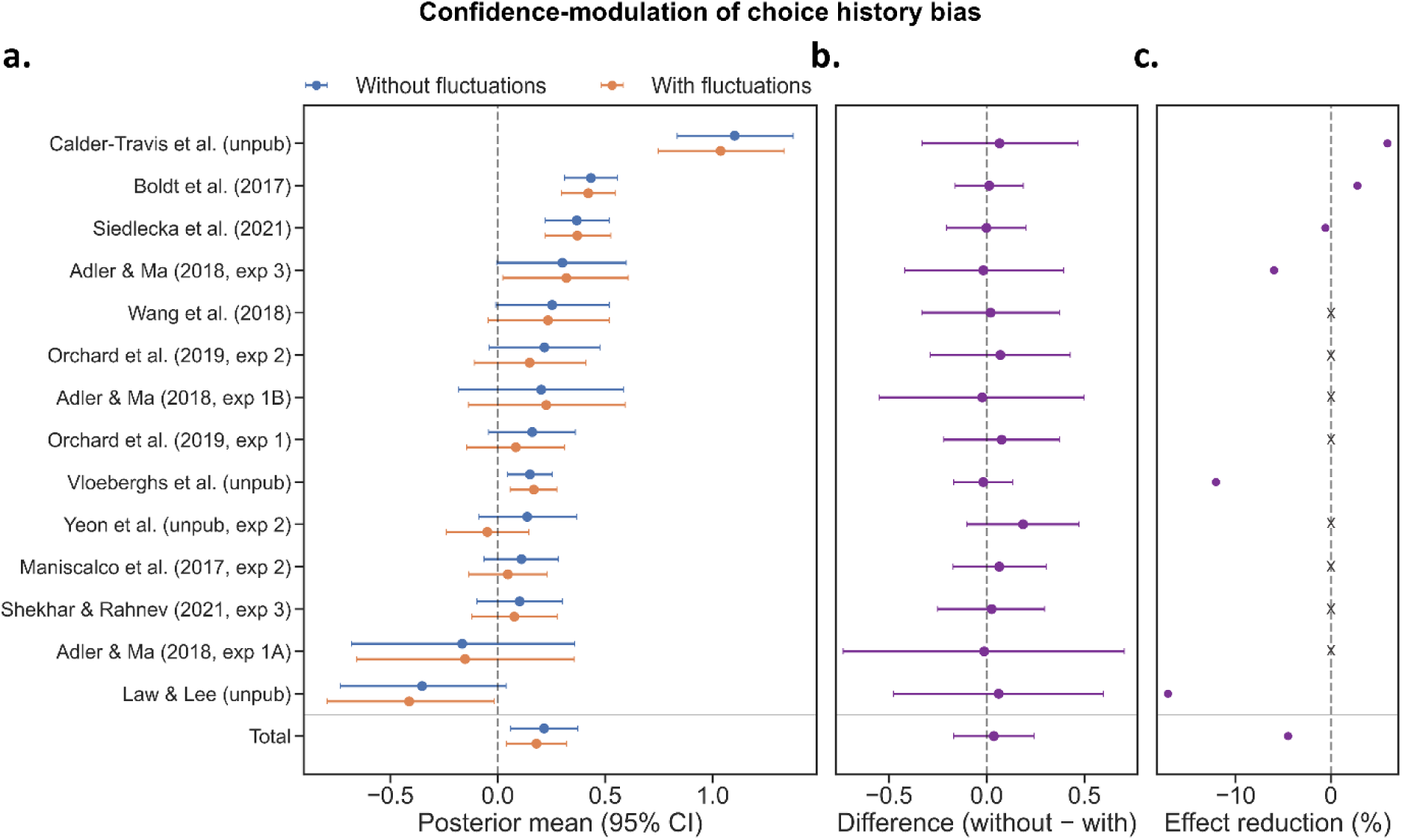
The confidence-modulation of choice history bias remains unaffected by criterion fluctuations. **A.** Posterior means and 95% credible interval for the interaction coefficient between previous response and previous confidence estimated with hMFC, with and without criterion fluctuations. In 4 out of 14 datasets the effect was significant without measuring criterion fluctuations. These effects remained in all four datasets after the inclusion of criterion fluctuations. Across datasets, we find a significant effect both before and after taking into account criterion fluctuations. **B.** None of the individual datasets showed a significant posterior difference, nor was there an effect across datasets. **C.** The four datasets showing a significant effect before accounting for criterion fluctuations exhibited virtually no change after criterion fluctuations were included, with an average reduction of 0.5% (range: −6.0% to 5.9%). Both datasets where the effect only emerged after including criterion fluctuations featured an average increase of 14.6%. Across all six datasets, accounting for criterion fluctuations increased the estimated effect by an average of 4.5%.To avoid interpreting percentage changes for effects that were not reliably present, this reduction metric was computed only for datasets in which the 95% credible interval of at least one of the two posterior estimates excluded zero.

The posterior differences between models further indicated that criterion fluctuations have little influence on the confidence modulation of choice history bias (Figure 6B). None of the individual datasets showed a significant change in the estimated effect after accounting for criterion fluctuations, nor was there evidence for an overall difference across datasets (*β_without−with_* = 0.036, CI(95) = [-0.17, 0.241]).

Lastly, the four datasets with a significant effect before accounting for criterion fluctuations showed an average reduction of 0.5% (range: −6.0 to 5.9) (Figure 6C). Both datasets where the effect emerged only after including criterion fluctuations featured an average increase of 14.6%. Across these six datasets, measuring criterion fluctuations resulted in an average increase of 4.5%. Together, these results suggest that criterion fluctuations do not explain the confidence-modulation of choice history bias.

## Discussion

Perceptual decision making is shaped not only by the current sensory input but also by experimental history. The influence of previous events, such as the preceding stimulus, the previous response, and the associated confidence, is generally referred to as sequential effects. Previous modelling work has generally assumed that experimental history exerts a *causal* influence on subsequent decisions. For example, within SDT, sequential effects can be explained by systematic shifts in the decision criterion as a function of the previous stimulus or response (Dyjas et al., 2012; Hachen et al., 2020; Lages & Treisman, 2010; Lee et al., 2023; Lee & Lee, 2023; Treisman & Williams, 1984). Similarly, Bayesian observer models account for sequential effects through the implementation of priors, which represent information from previous trials (Fritsche et al., 2020; Van Geert et al., 2024; Wei & Stocker, 2015). An alternative mechanism, fundamentally different from the previous models, proposes that sequential effects can be generated by trial-to-trial fluctuations in the decision criterion, *independent* of experimental history. For instance, Vloeberghs and colleagues (2025) showed that an attractive effect of previous response (i.e., a tendency to repeat; choice history bias) can be generated with an SDT criterion fluctuating from trial-to-trial according to an first-order autoregressive process (AR1). Crucially, these fluctuations were generated completely independent of experimental history, hereby questioning the assumed causal role, typically attributed in previous computational models (see also Gupta & Brody, 2022; Lak et al., 2020 for similar simulations).

In the current work, we extended these simulations by showing that criterion fluctuations can be a mechanism for a broader range of sequential effects, including the confidence modulation of choice history bias and the repulsive effect (i.e., tendency to alternate) of the previous stimulus. Using hMFC, we simultaneously estimated the influence of experimental history and criterion fluctuations. Across 14 empirical datasets, we found that the effect of both previous stimulus and previous response were strongly influenced by criterion fluctuations, while the confidence-modulation of choice history bias remained largely unaffected.

Across all datasets, the repulsive effect of the previous stimulus entirely disappeared after controlling for criterion fluctuations. At the individual dataset level, the effect became non-significant in 4 of the 10 datasets in which it was initially present, with an average reduction in effect size of 56.8%. Criterion fluctuations also substantially altered estimates of the previous-response effect (i.e., choice history bias), producing an overall shift towards alternation. All four datasets that initially showed a tendency to repeat became non-significant after accounting for criterion fluctuations. In three additional datasets, controlling for criterion fluctuations revealed a tendency to alternate that was absent in the original model, while in the two datasets that already showed a repulsive effect, this tendency became even stronger. In contrast, the confidence modulation of choice history bias was largely unaffected by criterion fluctuations. The effect remained significant across datasets after controlling for criterion fluctuations, and at the individual dataset level all four datasets with a significant effect remained significant.

Our findings indicate that criterion fluctuations account for a substantial portion of the effect of the previous stimulus, with the effect disappearing after controlling for these fluctuations in an analysis across all datasets. The mechanism underlying this sequential effect has long been debated. One account attributes it to sensory adaptation, whereby repeated stimulation reduces neuronal firing rates in early sensory areas such as V1 (Fritsche et al., 2017; Pascucci et al., 2019; Sadil et al., 2024). Extensions for Bayesian observer models have also been proposed to explain the repulsive effect of previous stimulus. For instance, efficient encoding allocates sensory resources according to the recent stimulus distribution, producing an asymmetric likelihood function with a long tail away from the most probable stimulus. This biases perceptual estimates away from recently encountered stimuli, giving rise to repulsive sequential effects (Fritsche et al., 2020; Van Geert et al., 2024; Wei & Stocker, 2015). A third account proposes that the decision criterion shifts toward the previous stimulus, increasing the likelihood of making the opposite response on the subsequent trial. Using neural recordings and computational modeling to dissociate the sensory adaptation account and the criterion shifting account, Lee and colleagues (2023) concluded that stimulus-induced criterion shifts best explained the repulsive effect. In contrast, our findings demonstrate that this effect is instead driven by criterion fluctuations *independent* of previous stimuli.

The current work also demonstrates that criterion fluctuations substantially influence estimates of choice history bias. The tendency to repeat the previous response has traditionally been attributed to observers’ internalized belief that the environment is temporally autocorrelated (Fischer & Whitney, 2014; Treisman & Williams, 1984; Yu & Cohen, 2008). Because the environment is assumed to be stable over short timescales, past events become informative about future ones. This idea has motivated models in which the decision criterion is updated based on previous responses, as well as standard Bayesian observer models in which posterior beliefs are systematically biased towards prior expectations derived from response history. In our analyses, we observed an attractive effect of the previous response in 4 of the 14 datasets. After accounting for criterion fluctuations, this effect disappeared in all four datasets. These findings suggest that attractive choice history biases are more likely to arise as a byproduct of criterion fluctuations than from an adaptive belief that the environment is autocorrelated. While the attractive effects are fully explained by criterion fluctuations, it remains an open question which mechanism then underlies the repulsive effect of previous response that remained in six datasets.

In contrast to the effects of the previous stimulus and previous response, criterion fluctuations had little to no impact on the confidence modulation of choice history bias (i.e., the interaction between previous response and previous confidence). One popular interpretation capitalizes on ideas from reinforcement learning, and proposes that confidence serves as an internal teaching signal in the absence of external feedback. According to this view, high-confidence choices generate a stronger reinforcement signal, strengthening the neural circuits that produced this decision (increasing the likelihood of repeating choices), whereas low-confidence choices generate weaker reinforcement, increasing the likelihood of switching on subsequent trials (Guggenmos et al., 2016; Lak et al., 2017, 2020). Bayesian observer models can also account for a part of this effect. High confidence on the previous trial corresponds to a relatively narrow posterior distribution. If this posterior is carried forward as the prior for the next trial, its greater precision increases its influence on the subsequent percept, thereby promoting repetition. However, it is less clear how these models explain the tendency to alternate following perceived-error trials. Because Bayesian updating shifts beliefs toward the prior, a symmetric likelihood function naturally predicts attractive effects. Without additional assumptions, the standard Bayesian framework is not capable of explaining the full range of the confidence-modulation of choice history bias.

While our work has direct implications for the computational mechanisms underlying sequential effects, it also highlights that criterion fluctuations could be a major confounding factor in the interpretation of empirical findings and associated theoretical work. For example, studies of autism spectrum disorder (ASD) and psychosis often use sequential effects as a probe of disrupted reliance on prior information, a characteristic frequently proposed to underlie these disorders (Bosch et al., 2022; Eckert et al., 2023; Feigin et al., 2021; Schaub et al., 2025). However, because criterion fluctuations can themselves generate sequential effects, stronger sequential effects in one group do not necessarily imply greater reliance on prior information. Instead, they may simply reflect larger fluctuations in the decision criterion. Consequently, without explicitly accounting for criterion fluctuations using hMFC it is difficult to draw valid conclusions about differences in the use of prior information between groups.

Criterion fluctuations may also change the interpretation of neural correlates of sequential effects. Numerous studies have investigated how neural signals encode previous stimuli and choices, and how this information is integrated into subsequent decision making. For example, Urai & Donner (2022) reported that previous responses were represented by gamma-band activity in the parietal cortex. Other studies have identified neural signatures of previous responses in the visual cortex (Lueckmann et al., 2018; St. John-Saaltink et al., 2016), although it remains debated whether these signals reflect feedforward or feedback processes (Macke & Nienborg, 2019). In addition, regions within the posterior parietal cortex (Akrami et al., 2018; Hwang et al., 2017, 2019; Scott et al., 2017) and prefrontal cortex (Barbosa et al., 2020; Mochol et al., 2021) have been implicated in sequential effects. However, if sequential effects are driven by fluctuations in decision criterion, as shown in our work, then some neural signals that appear to reflect the influence of preceding stimuli or responses on subsequent decisions may instead represent the decision criterion fluctuating over trials in an autocorrelated fashion. Therefore, researchers interested in the neural underpinnings of sequential effects should explicitly control for criterion fluctuations.

While the current work focuses on three sequential effects, criterion fluctuations can theoretically account for a much broader range of sequential effects. First, they can explain the exponentially decaying influence of more distant trial lags, whereby trials further in the past exert weaker effects on current behavior (Fritsche et al., 2020; Hachen et al., 2020; Urai et al., 2019). This follows naturally from the AR(1) structure of the criterion fluctuations, which itself is characterized by a decaying dependence on previous lags. Criterion fluctuations also provide an elegant explanation for individual differences in the strength of sequential effects. For example, Van Geert et al. (2022) found that the magnitude of different sequential effects was correlated within individuals: a stronger tendency to repeat previous responses was associated with a stronger tendency to alternate as a function of the previous stimulus. Criterion fluctuations can explain this phenomenon through a direct link between the magnitude of criterion fluctuations and the resulting sequential effects. Individuals with more pronounced fluctuations (i.e., higher values AR(1) parameters *ϕ* and/or *σ*) are expected to exhibit both stronger repetition and alternation tendencies. Finally, sequential effects have been shown to exhibit substantial temporal stability across sessions, with both the direction and magnitude of these biases remaining highly correlated over time (Van Geert et al., 2022). If the factors underlying criterion fluctuations represent stable individual traits, this could explain why individuals who display strong criterion fluctuations at one point in time continue to exhibit similarly strong fluctuations at later points. However, this last point is highly speculative given that it is currently unclear which factors underlie criterion fluctuations.

In conclusion, in the current work we disentangled causal contributions to experimental history, typically assumed in computational models of sequential effects, from non-causal contributions to sequential effects induced by fluctuations in the SDT decision criterion. Across 14 empirical datasets, criterion fluctuations fully explained the repulsive effect of the previous stimulus and substantially altered estimates of choice history bias. These findings suggest that a substantial portion of sequential dependencies previously attributed to learning or adaptation may instead reflect endogenous fluctuations in decision criterion, highlighting the need to account for these fluctuations when studying the computational mechanisms underlying sequential effects.

---

**Supplementary Table 1:**
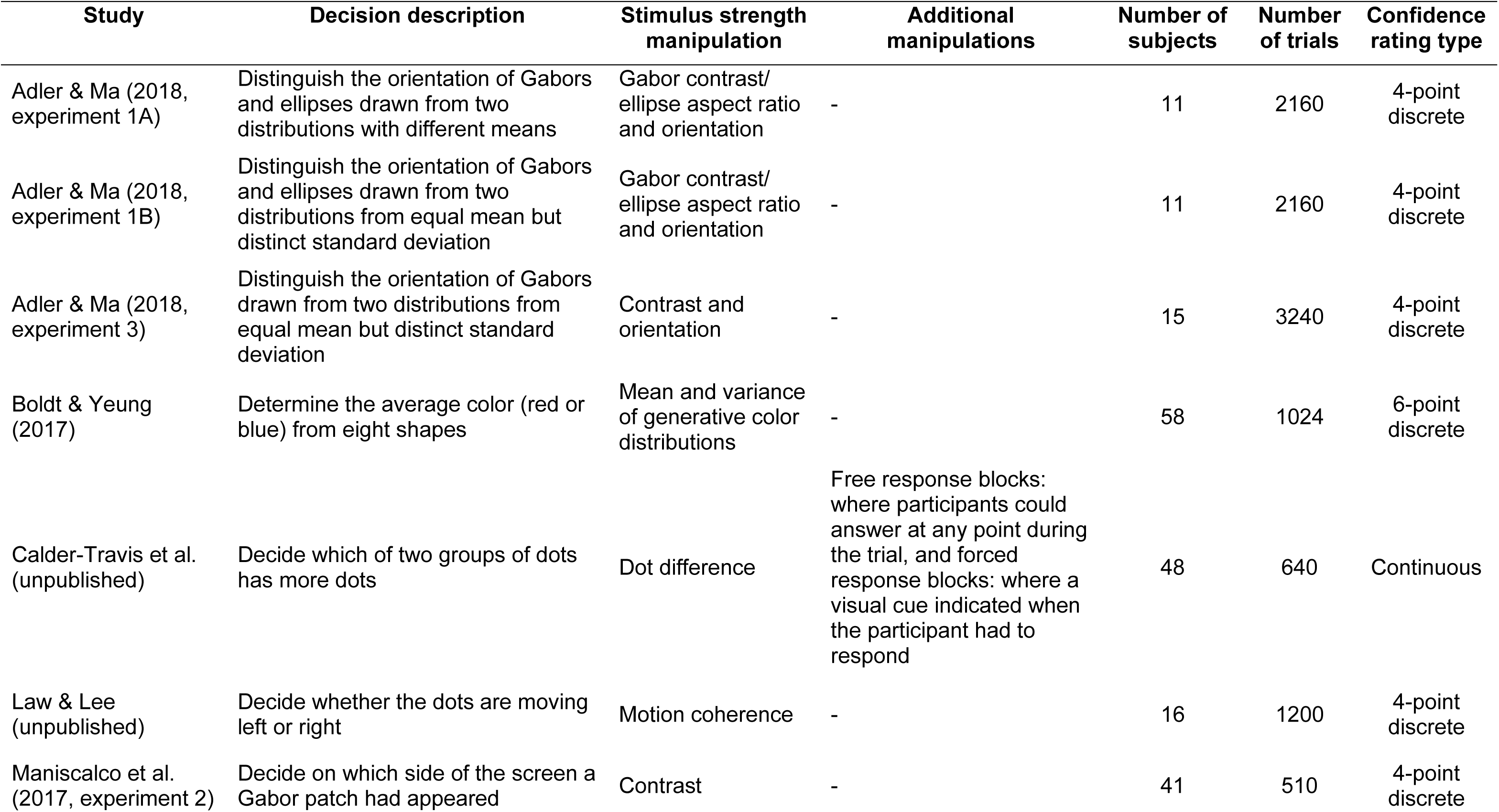
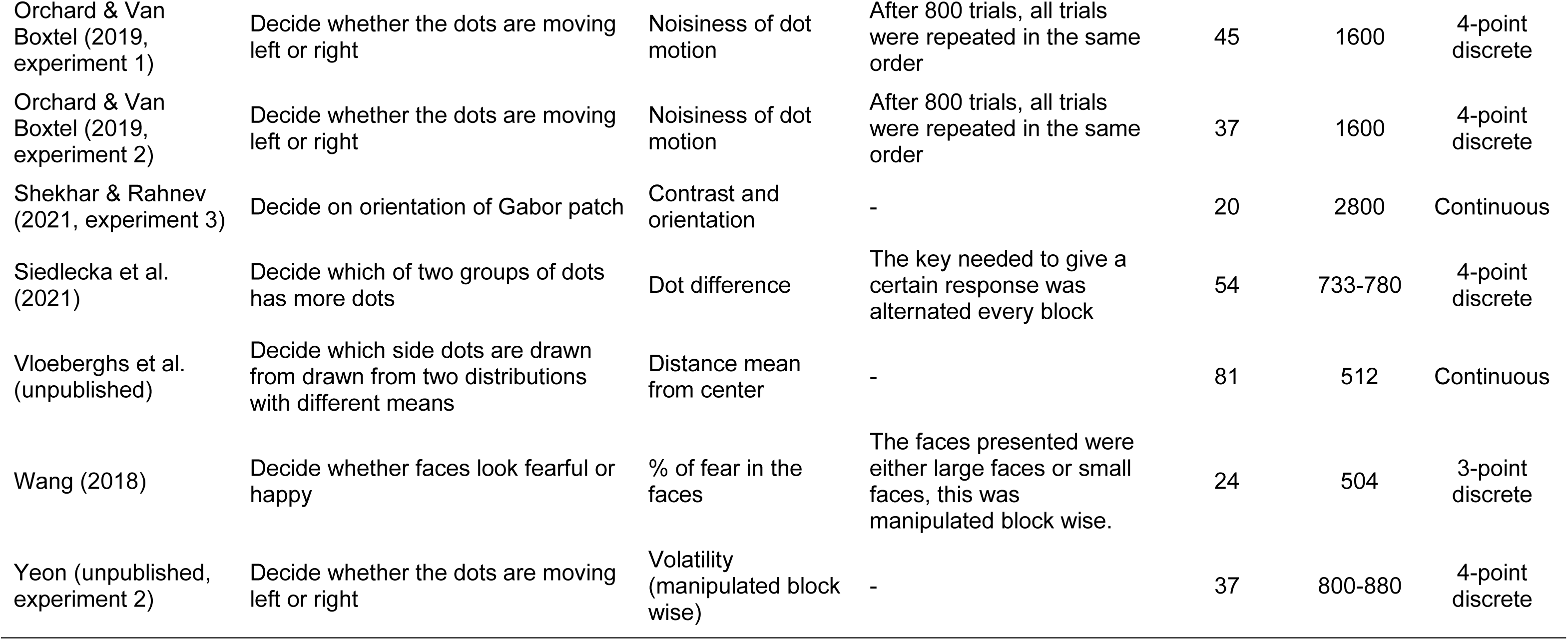
Detailed description of empirical datasets.

## Notes

### Competing Interest Statement

The authors have declared no competing interest.

